# Linking power, efficiency, and bifurcations in consumer–resource systems

**DOI:** 10.1101/2025.07.14.664758

**Authors:** Serguei Saavedra, Yuguang Yang, Christopher P. Kempes, Chengyi Long, Ricard Solé, Tomo Yoshino, Marco Tulio Angulo

## Abstract

Three hypotheses help organize energetic thinking about living systems: Lotka’s *Maximum Power Principle*, Odum–Pinkerton’s *Intermediate Efficiency Principle*, and Morowitz’s *Biological Cycling Principle*. Here we show how these hypotheses fit together in consumer–resource systems, moving from qualitative principles to formal, testable statements. Using the Rosen-zweig–MacArthur model, we prove that the consumer’s maximum output power lies exactly on the Hopf boundary that separates stable points from cycles; at that point, the resulting power efficiency is intermediate. In the Rosenzweig–MacArthur model the Hopf is supercritical, so a stable limit cycle appears smoothly as the equilibrium loses stability. We treat the Hopf onset of time-periodic population oscillations as a population-level analogue of sustained cycling under energy flux. We then embed these energetic statements in adaptive dynamics: with convex trait costs, evolutionary singular strategies exist and are locally stable, but they coincide with the maximum-power state only under explicit marginal-cost conditions. Together, these results unify classic ideas in the concrete setting of consumer-resource systems and suggest measurements to evaluate when bifurcations, energetics, and evolution can converge.

## Introduction

Ecological systems, ranging from microbiomes to rainforests, are among the most complex, living, dissipative systems in nature (de Castro and McShea, 2022, Murphy and O’Neill, 1995, Odum, 1983). For example, energy from the Sun is first converted by plants, then transferred to herbivores that feed on plants, and subsequently to carnivores that feed on herbivores, continuing through the interconnected web of life (Odum and Barrett, 2005, Olff et al., 2009, Pielou, 2001). This ecological complexity has been addressed over the years following different theoretical approaches (Solé and Bascompte, 2005). In most cases, nonlinear dynamical systems have provided a solid framework for exploring the nature of ecological attractors at the population level (Case, 1999, Strogatz, 2014). However, ecological systems are also open systems, where energy flows (such as the input of solar radiation and the dissipation of metabolic heat) govern processes of self-organization, trophic interactions, and long-term stability (Svirezhev, 2000). Both sides of the coin have been treated in the context of a system view of complexity, where out-of-equilibrium phenomena appear to be described in terms of bifurcations and phase transitions (Haken, 1987, Nicolis and Prigogine, 1989). How are nonlinear dynamics and thermodynamics related to each other in the context of ecology and evolution?

Consumer–resource nteractions are the workhorses of population ecology (MacArthur, 1955). Because they are open subsystems that take in energy and dissipate it as heat, they also offer a natural stage on which to examine how energetic constraints and nonlinear dynamics meet (Lindeman, 1942). Our aim is to revisit three classic hypotheses—Lotka’s *Maximum Power*, Odum–Pinkerton’s *Intermediate Efficiency*, and Morowitz’s *cycling theorem*—and to show, within consumer-resource systems, how they connect and where their limits are. In the 1920s, Lotka Lotka (1922), building on Boltzmann, proposed what became known as the *Maximum Power Principle*: among competing systems, configurations that capture and transform *usable* energy at a higher rate (power) are favored by selection. In the 1950s, Odum and Pinkerton (Odum and Pinkerton, 1955) refined this idea by showing—via network analogies and linear nonequilibrium thermodynamics—that maximum power is achieved at intermediate conversion efficiency, not at perfect efficiency; this is the *Intermediate Efficiency Principle* (akin to source–load matching in circuits). In parallel, Morowitz (Morowitz, 1966) established a complementary, flux-based result: in driven nonequilibrium networks at steady state, continuous energy throughput necessarily produces material cycling—flows around closed loops (often called the cycling theorem, here referenced as the *Biological Cycle Principle*). In our setting, Lotka and Odum–Pinkerton can be interpreted as *selective* influences that filter which ecological attractors persist (by balancing power and efficiency), whereas Morowitz provides a *generative* constraint: sustained energy flux organizes systems that exhibit persistent cycling of matter, thereby shaping the ecological states on which selection can act (Levin, 1998, Saavedra, 2024, Solé et al., 2024).

A central contribution here is to turn these hypotheses into formal theorems in a widely used CR model, with proofs in the Appendix and plain-language statements in the main text. Our hope is not to elevate one model above all others, but to demonstrate that energetic concepts can be made exact in a way that invites testing. To keep the discussion concrete, we work with the Rosenzweig–MacArthur model, which captures a self-regulated resource consumed with a Holling type II response (Rosenzweig and MacArthur, 1963). We define power in familiar biological units—biomass times specific growth—so that input power corresponds to primary production and output power to assimilated consumer production. This choice makes contact with chemostat and microcosm measurements. We then show that the trait value that maximizes consumer output power lies precisely on the Hopf boundary, and that at this point the system’s power efficiency is necessarily intermediate. In the Rosenzweig–MacArthur model, the Hopf is supercritical, so the transition from a stable point to a stable cycle is continuous.

Morowitz’s contribution plays a complementary role. His Biological Cycling Principle establishes that, at steady state, driven reaction networks necessarily exhibit material cycling—that is, there are flows around closed loops in the network (Morowitz, 1966). This is a statement about the topology of fluxes at steady state, not about time-periodic oscillations. In the population model, by contrast, cycles arise dynamically via a Hopf bifurcation as assimilation increases. We therefore read the Hopf threshold as a population-level analogue of Morowitz-style sustained cycling under energy flux: both statements identify when continuous energy input organizes the system into cycling regimes—one in network space, the other in time.

Finally, we ask how evolution positions consumer-resource systems on this energetic map. We do not assume that natural selection maximizes a physical quantity (such as power). Instead, using adaptive dynamics, we derive selection gradients for foraging effort and assimilation efficiency under convex mortality costs (Doebeli, 2011). We show that evolutionary singular strategies are locally stable and identify explicit testable conditions under which these singular points align with the maximum-power state. The alignment depends on marginal costs and enrichment; outside those conditions, selection systematically departs from maximum power even as energy flow continues to structure the feasible regimes.

### Model

We study the Rosenzweig–MacArthur consumer–resource model with a Holling type II response:

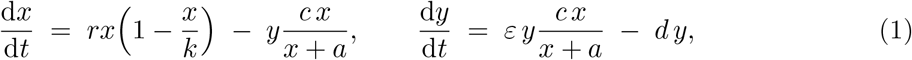

where *x* and *y* are resource and consumer biomasses. Parameters *r* (intrinsic resource growth) and *d* (mortality) have units [time^−1^]; *k* (carrying capacity) and *a* (half-saturation biomass) have units [mass]; *c* (max per-biomass consumption) has units [time^−1^]; *ε* ∈ [0, 1] is assimilation efficiency. We assume *k > a* and interpret *r* and *k* phenomenologically as the net environmental balance of uptake, assimilation, and crowding losses. The Rosenzweig–MacArthur model provides a minimal, biologically interpretable stage where intake saturation (*a*), enrichment (*k*), and assimilated intake (*cε*) can be linked to feasibility, energetics, and adaptive dynamics.

### Bifurcation

In the context of the Biological Cycle Principle (Morowitz, 1966), the coupled consumer–resource system of Eq. (1) can exhibit either an asymptotically stable equilibrium at (*x*^*^, *y*^*^), or a stable limit cycle around this equilibrium that emerges through a Hopf bifurcation (Strogatz, 2014). Solving Eq. (1) for interior equilibria yields

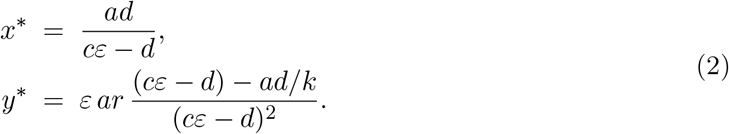

Feasibility (*x*^*^ *>* 0, *y*^*^ *>* 0) requires

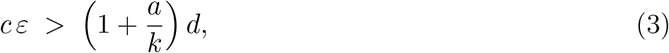

i.e., the maximum assimilated intake *cε* must exceed mortality by a buffer (*a/k*)*d* due to saturation (*a*) and finite capacity (*k*).

Linearizing at (*x*^*^, *y*^*^) shows a Hopf bifurcation when the Jacobian trace vanishes, yielding

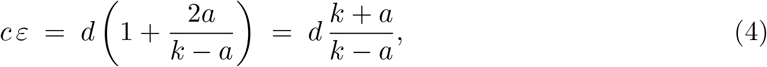

which strictly exceeds Eq. (3) for *k > a*. In the RM model this Hopf is *supercritical* (Strogatz, 2014): below Eq. (4) the interior point is asymptotically stable; at equality, a Hopf occurs; just above, a small stable limit cycle emerges (Appendix). The qualitative regimes partitioned by Eqs. (3) and (4) are visualized in Appendix (for an *ε*-scan at fixed *c*).

In other words, oscillations arise only when the consumer assimilates energy at a rate high enough to overcome both mortality *d* and the damping effects 2*ad/*(*k* −*a*) introduced by resource saturation and limited carrying capacity. By rewriting 2*ad/*(*k* − *a*) = (*ad/k*) [2*k/*(*k* − *a*)], we note that 2*k/*(*k* − *a*) is decreasing in *k* for *k > a* and 2*k/*(*k* − *a*) *>* 2. Therefore, oscillations naturally require feasibility since 2*ad/*(*k* − *a*) ≥ 2*ad/k > ad/k*.

### Energetics

Following Odum and Pinkerton (1955), we define input and output power as

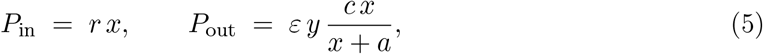

i.e., power is measured as biomass times specific growth (a flux). In Eq. (1), *P*_in_ is the rate at which energy enters via resource production (primary production), whereas *P*_out_ is the rate of assimilated consumer production. This formalism is appropriate for biotic systems and differs from classical abiotic chemical setups (Kempes et al., 2021). Dissipation occurs through (i) crowding losses in the resource (−*rx*^2^*/k*), (ii) consumer mortality (−*dy*), and (iii) incomplete assimilation (the fraction 1 − *ε* of extracted energy). Other power conventions are possible; we adopt Eq. (5) because they map cleanly to ecological measurements.

Long-term averages (exact at equilibria; period averages on cycles) yield (Appendix)

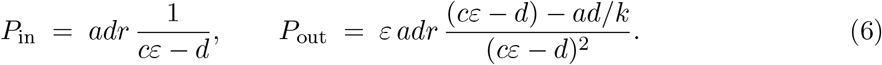

Hence the power efficiency

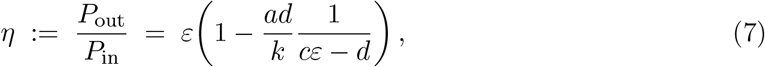

so *η < ε* whenever feasibility (Eq. (3)) holds: finite *k* and saturation *a* impose structural losses. This power efficiency is distinct from the consumer’s capture/assimilation efficiency; bundling *b* = *cε* would trivially give *η* = *b/c* = *ε*, but that definition omits the structural losses explicitly captured in Eq. (7).

At equilibrium, output power depends only on the standing resource:

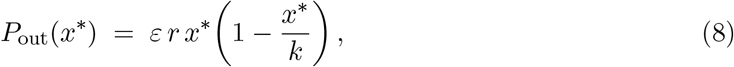

a concave quadratic maximized at *x*^*^ = *k/*2 (Appendix). This state-function view is a practical diagnostic: systems near maximum power should sit at *x* ≈ *k/*2.

Maximizing *P*_out_ with respect to a single trait gives two complementary results. (i) Vary *c* at fixed *ε*:

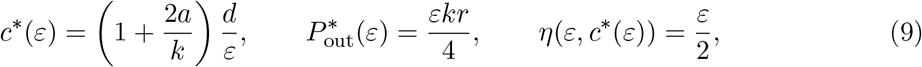

with *c*^*^*ε* = (2*a* + *k*)*d* /*k*, which lies below the Hopf threshold (Eq. (4)) when *k > a*. Thus, optimizing *c* yields maximum power at intermediate efficiency and a stable equilibrium (pre-Hopf).

(ii) Vary *ε* at fixed *c*:

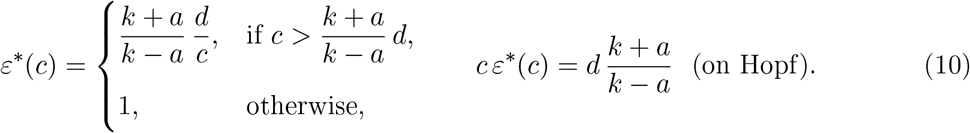

Here the power maximum lies exactly on the Hopf boundary: increasing *ε* to *ε*^*^(*c*) places the system at the onset of oscillations. At this Hopf point the power efficiency is *η*_Hopf_ = *ε* (*k* + *a*) /(2*k*), which lies strictly between *ε/*2 and *ε* for *k > a* (whereas *η* = *ε/*2 holds at the *c*-optimum; see Eq. (9)). Fig. 1a illustrates these statements for an *ε*-scan (fixed *c*): as *ε* increases, the system goes from infeasible to a stable point and then crosses the Hopf line; the output power peaks precisely at that boundary, where cycles emerge. In this sense, *ε*^*^(*c*) plays a “triple point” role: maximum power, intermediate efficiency, and the onset of oscillations.

**Figure 1.**
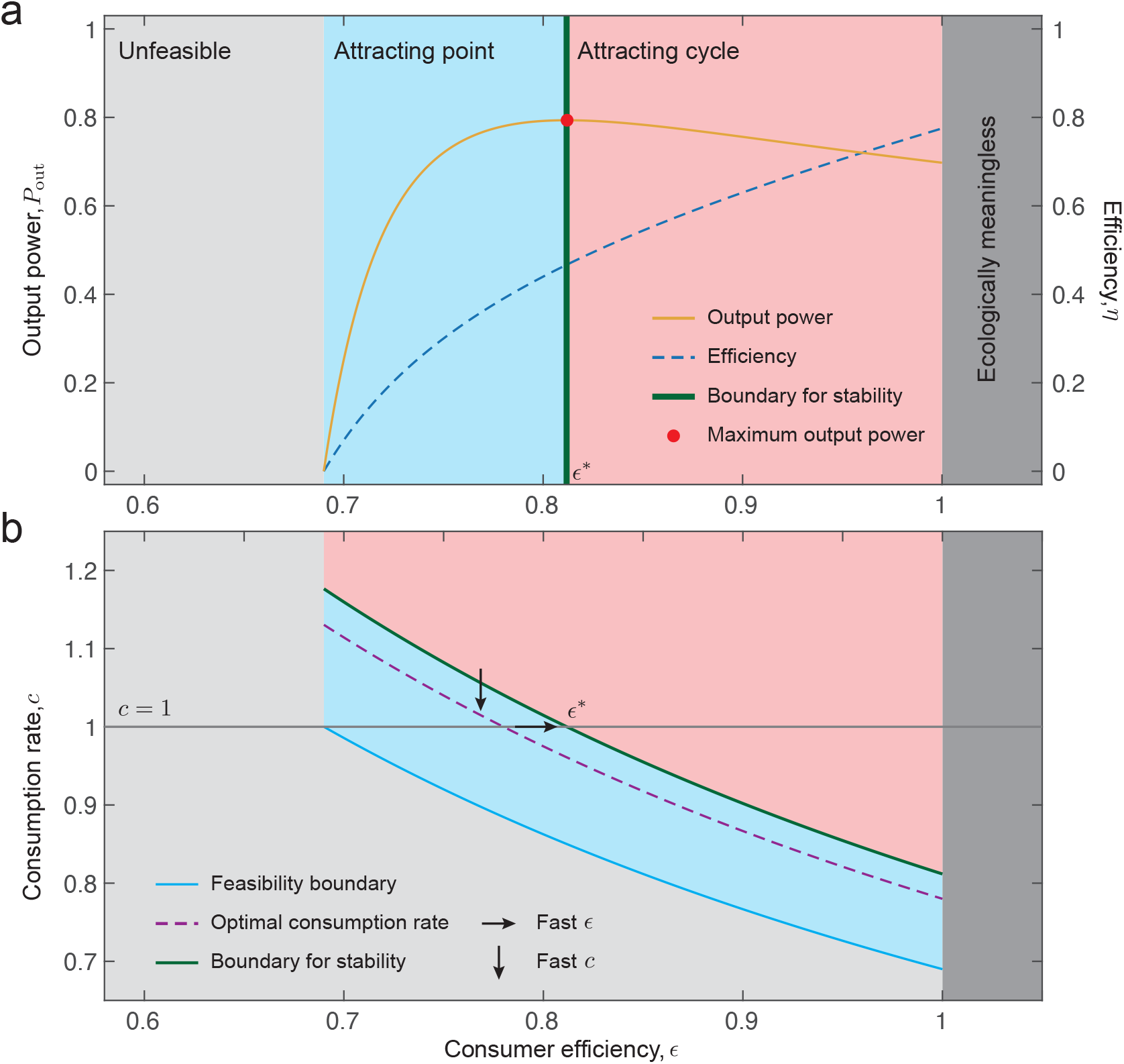
Bifurcations, energetics, and evolution. Numerical example (Supplementary Information). **(a)** Output power and efficiency analysis. As the consumer efficiency *ε* increases, the system’s efficiency increases (blue dashed line), while the output power (yellow line) first increases and then decreases. Colored regions represent distinct system behaviors: the left-most light-gray region is unfeasible, the blue region corresponds to an attracting fixed point (i.e., local asymptotic stability), the red region indicates an attracting limit cycle, and the dark gray region is biophysically impossible (i.e., *ε >* 1). The green vertical line marks the boundary between the blue and red regions (Hopf threshold, Eq. (4))). The red dot denotes the point of maximum output power, which occurs at the green boundary; the corresponding assimilation (consumption) efficiency is denoted by *ε*^*^. **(b)** Corresponding (*ε, c*)-plane illustrating the feasibility boundary (blue line), the stability boundary (green line), and the optimal consumption rate (purple dashed line). The gray horizontal line indicates *c* = 1, consistent with panel (a). Arrows denote fast adaptive directions: *fast ε* evolution (horizontal) drives the system toward the Hopf boundary, while *fast c* evolution (vertical) moves it toward the optimal consumption curve.

### Adaptive dynamics

We can draw evolutionary insights by treating assimilation efficiency *ε* and consumption rate *c* as adaptive traits of the consumer. Because *ε c*^*^(*ε*) ≠ *c ε*^*^(*c*), no single pair (*ε, c*) simultaneously maximizes output power for both traits—only in the limit *k* → ∞ do both best responses coincide (in line with classical chemical systems (Kempes et al., 2021)). When traits evolve on different timescales, the faster trait reaches its adaptive optimum while the slower one tracks a best response (Cortez and Ellner, 2010, Hofbauer and Sigmund, 1998).

We analyze one evolving trait at a time with convex mortality costs: *d*_*c*_(*c*) for effort and *d*_*ε*_(*ε*) for efficiency. The predator zero-growth condition at the interior equilibrium is

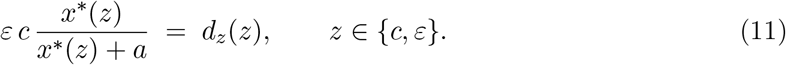

For effort, invasion fitness and selection gradient are

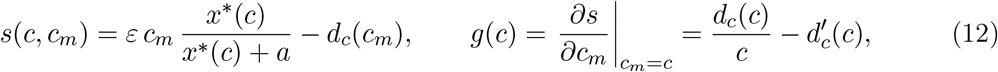

so the singular condition is

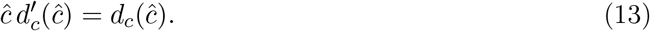

For efficiency,

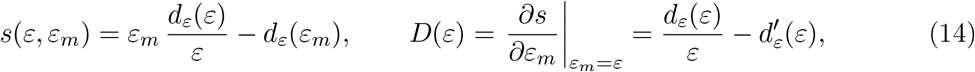

with singular

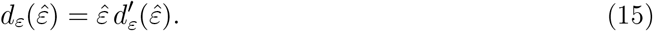

Under strict convexity 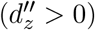, both singular points are ESS and convergence stable (CSS).

Because *P*_out_(*x*) is maximized at *x* = *k/*2 (Eq. 8), alignment between the CSS and maximum power occurs iff

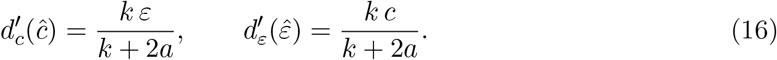

Outside these marginal-cost conditions, selection extremizes *d*_*c*_(*c*)*/c* or *d*_*ε*_(*ε*)*/ε* and generally does not maximize *P*_out_.

Fig. 1b illustrates these statements. If one trait adapts faster, it approaches its CSS while the other tracks a best response. Fast-*ε* dynamics carry the system to the Hopf “triple point” (maximum power, intermediate efficiency, onset of cycles via Eq. 10); fast-*c* dynamics approach a “double point” *c*^*^(*ε*) (Eq. 9), delivering maximum power at intermediate efficiency but pre-Hopf. The faster trait determines whether systems settle at the Hopf boundary (fast *ε*) or stay pre-Hopf (fast *c*).

## Discussion

Our goal was to connect three classic energetic ideas—Lotka’s *Maximum Power*, Odum–Pinkerton’s *Intermediate Efficiency*, and Morowitz’s *Biological Cycling* —to the concrete mechanics of consumer–resource systems, and to move from qualitative principles to formal testable theorems. Within the Rosenzweig–MacArthur model we established: (i) a feasibility bound (Eq. 3) that ties coexistence to the maximum assimilated intake *cε*; (ii) a supercritical Hopf threshold (Eq. 4) that cleanly separates fixed points from cycles; (iii) a “which-trait-varies” resolution of maximum power—on the pre-Hopf side when optimizing over effort *c* (with *η* = *ε/*2; Eq. 9) and exactly on the Hopf boundary when optimizing over *ε* (Eq. 10); (iv) a structural bound *η < ε* (Eq. 7) due to finite carrying capacity and intake saturation; and (v) a state-function view of production, *P*_out_(*x*^*^) = *εrx*^*^(1 − *x*^*^*/k*) (Eq. 8), maximizing at *x*^*^ = *k/*2 and extending to small cycles by averaging. Together, these statements turn a trio of long-standing principles into an energetic map for CR dynamics that can be checked directly.

Morowitz’s Biological Cycling Principle is a steady-state, flux-topology statement: continuous energy throughput in driven networks necessarily produces material cycling (flows around closed loops). Our population-level cycles arise dynamically via a supercritical Hopf as *cε* or *k* increase (Eq. 4). We therefore treat the Hopf onset as a population analogue of sustained cycling under energy flux: Morowitz identifies where fluxes must circulate in network space, whereas the Rosenzweig–MacArthur model shows when time-periodic cycles emerge in phase space as assimilation/enrichment rise. Clarifying this distinction prevents conflating steady-state flux cycling with temporal oscillations while still revealing a common energetic driver.

Casting power and efficiency in biological units (Eq. 5) and proving the bounds and coincidences above makes the principles falsifiable. For instance, Odum–Pinkerton’s “intermediate efficiency” becomes *η* = *ε/*2 at the *c*-optimum (Eq. 9), and Lotka’s “maximize power” becomes a precise location—either on the Hopf boundary when *ε* varies (Eq. 10) or pre-Hopf when *c* varies. The state-function *P*_out_(*x*^*^) (Eq. 8) yields a practical diagnostic: empirical systems near maximum power should sit at *x* ≈ *k/*2.

Using adaptive dynamics with convex trade-off costs, we derived selection gradients and showed singular strategies are CSS for both effort and efficiency (Eqs. 12–15). Crucially, evolved states align with maximum power only under explicit marginal-cost conditions (Eq. 16) that tie trait costs to enrichment and handling. Outside those conditions, selection extremizes *d*_*z*_(*z*)*/z* rather than *P*_out_, so one should not expect an optimization principle for power in general. Timescale separation clarifies outcomes: faster evolution in *ε* steers systems to the Hopf “triple point” (maximum power, intermediate efficiency, onset of cycles), whereas faster evolution in *c* approaches a “double point” with maximum power at intermediate efficiency but pre-Hopf stability.

Chemostats and plankton microcosms can be natural testbeds. Vary enrichment (*k*) while tracking (*x, y*) and estimating (*r, c, a, ε, d*); verify the Hopf relation *cε* = *d*(*k* + *a*)*/*(*k* −*a*) (Eq. 4); check that the production peak occurs at Hopf when *ε* varies but remains pre-Hopf when *c* varies; and use *x* ≈ *k/*2 (Eq. 8) as a parameter-light indicator of proximity to maximum power. Short-term selection or plasticity experiments that tilt the faster trait (effort vs. efficiency) can discriminate “triple” vs. “double” points. Finally, measure the local slope of mortality–trait trade-offs to test the alignment conditions in Eq. 16. If the goal is high production with predictable dynamics, tuning enrichment (*k*) and interaction mechanics (*a*) can place systems near *x* = *k/*2 without crossing the Hopf boundary, whereas indiscriminately pushing efficiency upward risks moving the system onto the Hopf line (Eq. 10) and into cycles.

Our theorems hold for consumer-resource systems with a self-regulated resource, a Type-II response, one evolving trait at a time, and convex mortality costs. We do not claim general optimization by natural selection. In richer settings—multiple species/resources, alternative functional responses, explicit stoichiometry, space, stochasticity, coevolution, non-convex trade-offs, or genetic constraints—optimization principles can fail and new phenomena (e.g., branching) can emerge. We view the results here as structural guides: (i) the coincidence of the power peak with the Hopf boundary when efficiency varies, (ii) intermediate power efficiency at the *c*-optimum, (iii) the *η < ε* bound, and (iv) explicit alignment conditions that connect marginal costs to enrichment and handling. By making the links between power, efficiency, bifurcations, and evolution explicit, we sharpen classic principles into concrete testable statements for consumer–resource systems, with measurable conditions to know when energetics and evolution can converge.

## Acknowledgments

SS acknowledges support from the National Science Foundation under Grant No. DEB-2436069. R.S. thanks the support of the Generalitat de Catalunya under grant AGAUR 2021 SGR 00751. MTA acknowledges support from UNAM-PAPIIT IA102225.

## —ONLINE APPENDIX—

### 1. Equilibria and feasibility analysis

The consumer–resource system studied is

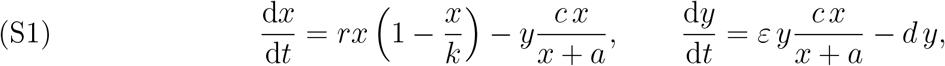

where *x* and *y* denote the biomasses of the resource and consumer, respectively (units [mass]). Parameters *r* and *k* are the resource regeneration rate (units [time]^−1^) and carrying capacity (units [mass]); the interaction is Holling Type II with consumption rate *c* (units [time]^−1^) and half-saturation *a* (units [mass]). The assimilation efficiency is *ε* ∈ [0, 1) (dimensionless) and *d* is mortality (units [time]^−1^).

The interior equilibrium (*x*^*^, *y*^*^) solves 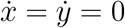:

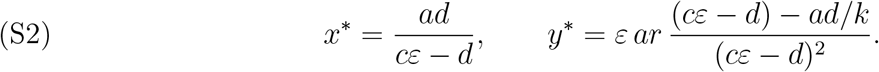

Feasibility (*x*^*^ *>* 0, *y*^*^ *>* 0) requires

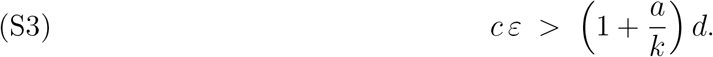

For numerical illustrations we use parameter values plausible for a lynx–hare system (Hudson Bay).

**Table 1.** Model parameters (biomass units: kg km^−2^ for *x, y*; yr^−1^ for rates).

| Symbol | Meaning (units) | Plausible range | Working value |
| --- | --- | --- | --- |
| $r$ | Intrinsic resource growth rate (yr<sup>−1</sup>) | 1.2–1.8 | 1.6 |
| $k$ | Carrying capacity (kg km<sup>−2</sup>) | 850–1200 | 1000 |
| $c$ | Per-lynx max intake (yr<sup>−1</sup>) | 50–70 | 60 |
| $a$ | Half-saturation prey biomass (kg km<sup>−2</sup>) | 40–80 | 60 |
| $\varepsilon$ | Assimilation efficiency (–) | 0.10–0.25 | 0.15 |
| $d$ | Adult lynx mortality (yr<sup>−1</sup>) | 0.20–0.45 | 0.30 |

### 2. Hopf bifurcation

A limit cycle appears via a Hopf bifurcation when the Jacobian *J* of (S1) at (S2) has purely imaginary eigenvalues, i.e., when trace(*J* ) = 0 with det(*J* ) *>* 0. For the Rosenzweig–MacArthur system,

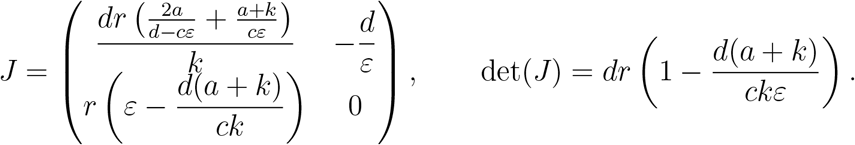

Under feasibility (S3), det(*J* ) *>* 0. The trace condition trace(*J* ) = 0 is equivalent to

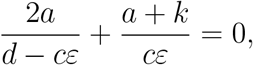

which yields the Hopf threshold

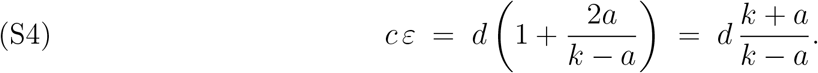

For *k > a*, (S4) strictly exceeds the feasibility bound (S3), so feasibility precedes the Hopf onset. In the RM model this Hopf is supercritical: below (S4) the interior equilibrium is asymptotically stable; at equality, a Hopf occurs; just above, a small-amplitude stable limit cycle emerges.

**Figure S1.**
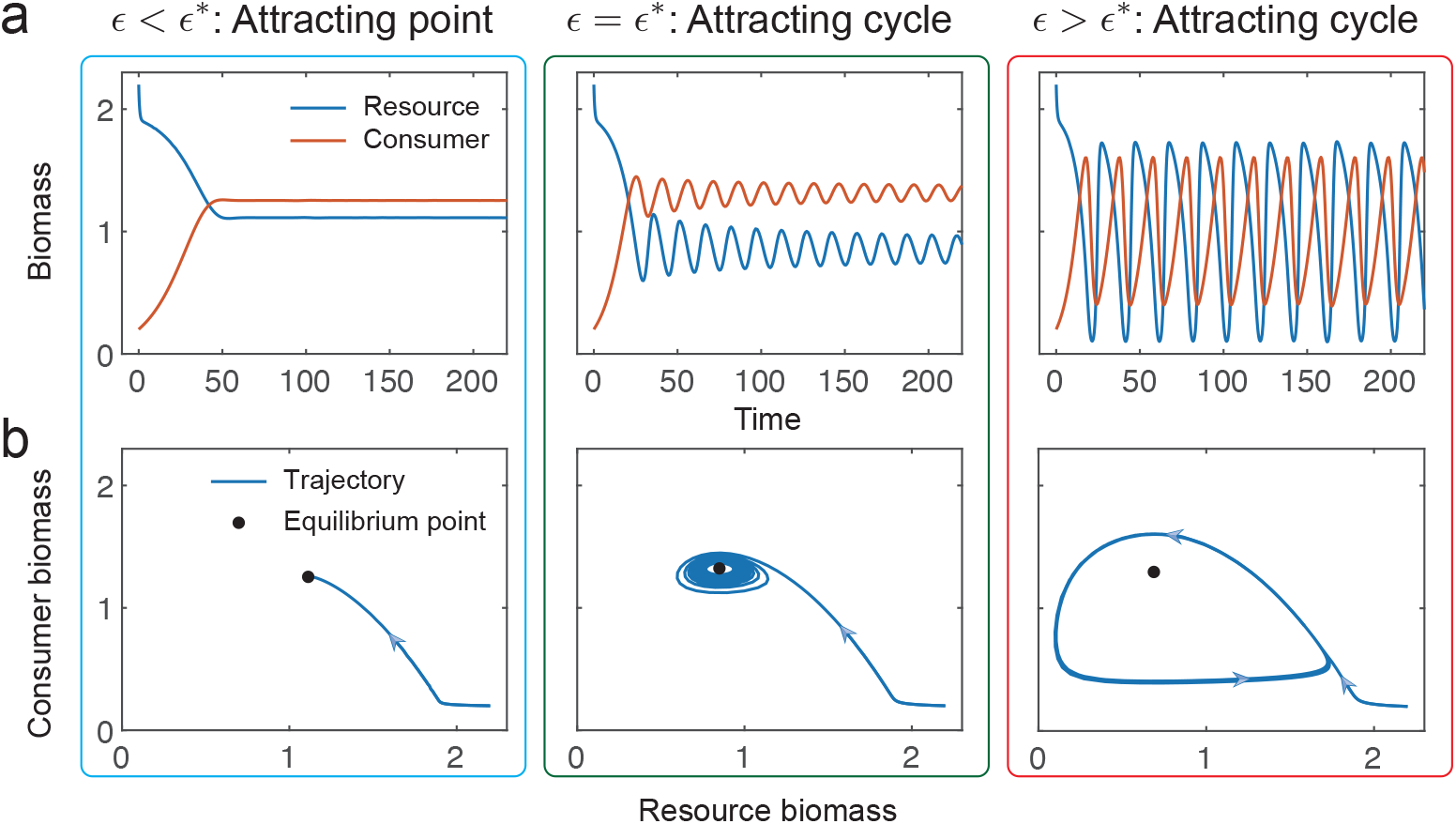
System dynamics as consumer efficiency *ε* increases. (**a**) Time series of resource and consumer biomasses. (**b**) Corresponding phase portraits. Left (blue): convergence to a stable equilibrium for *ε < ε*^*^. Middle (green): transition at *ε* = *ε*^*^ (Hopf onset; cf. (S4)). Right (red): sustained cycles for *ε > ε*^*^.

### 3. Long-term average power

For an integrable function *g*(*t*), define the long-term average

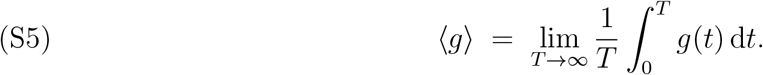

When (S1) is feasible, trajectories either converge to the interior equilibrium or settle onto a limit cycle born at the Hopf bifurcation. In either case (equilibrium as a degenerate 0-period orbit), the average of every time derivative is zero; e.g.,

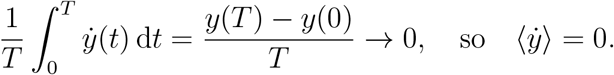

Following the main text, define

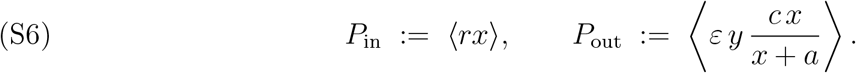

Averaging (S1) gives

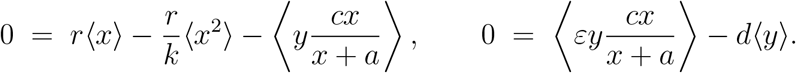

Hence,

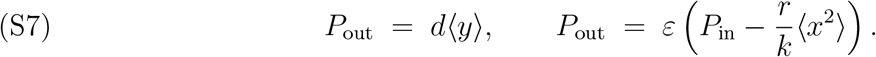

In the Volterra limit (Type I, *k* → ∞) time averages equal equilibrium values [1, pp. 16]. For the present Type II model, these identities remain exact at a stable interior equilibrium; substituting (S2) yields the closed forms

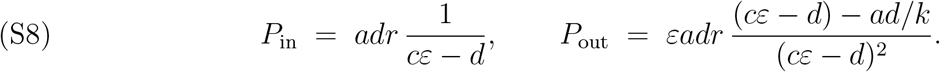

Define the *power efficiency*

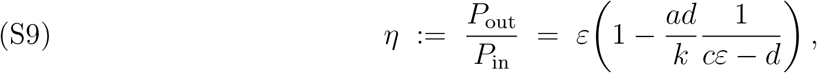

so *η < ε* whenever (S3) holds.

#### Accuracy of equilibrium formulas beyond Hopf

Let *µ* := *cε* − (*cε*)_*H*_ with 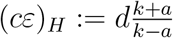 from (S4). At a supercritical Hopf, the centre-manifold normal form

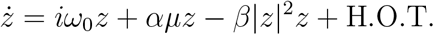

implies the limit-cycle amplitude 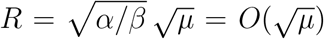. Writing deviations (*u, v*) = (*x* − *x*^*^, *y* − *y*^*^) and transforming to 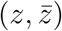 shows ⟨*u*⟩ = ⟨*v*⟩ = 0 to *O*(*R*) and that quadratic averages contribute only through *r*⟨*x*⟩ − (*r/k*)⟨*x*^2^⟩, whose *O*(*µ*) term cancels at Hopf because *x*^*^ = (*k* − *a*)*/*2. Hence

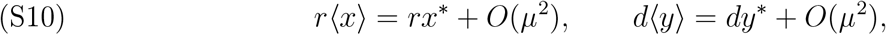

so (S8) approximates period-averaged powers with *O*(*µ*^2^) error just beyond Hopf.

**Figure S2.**
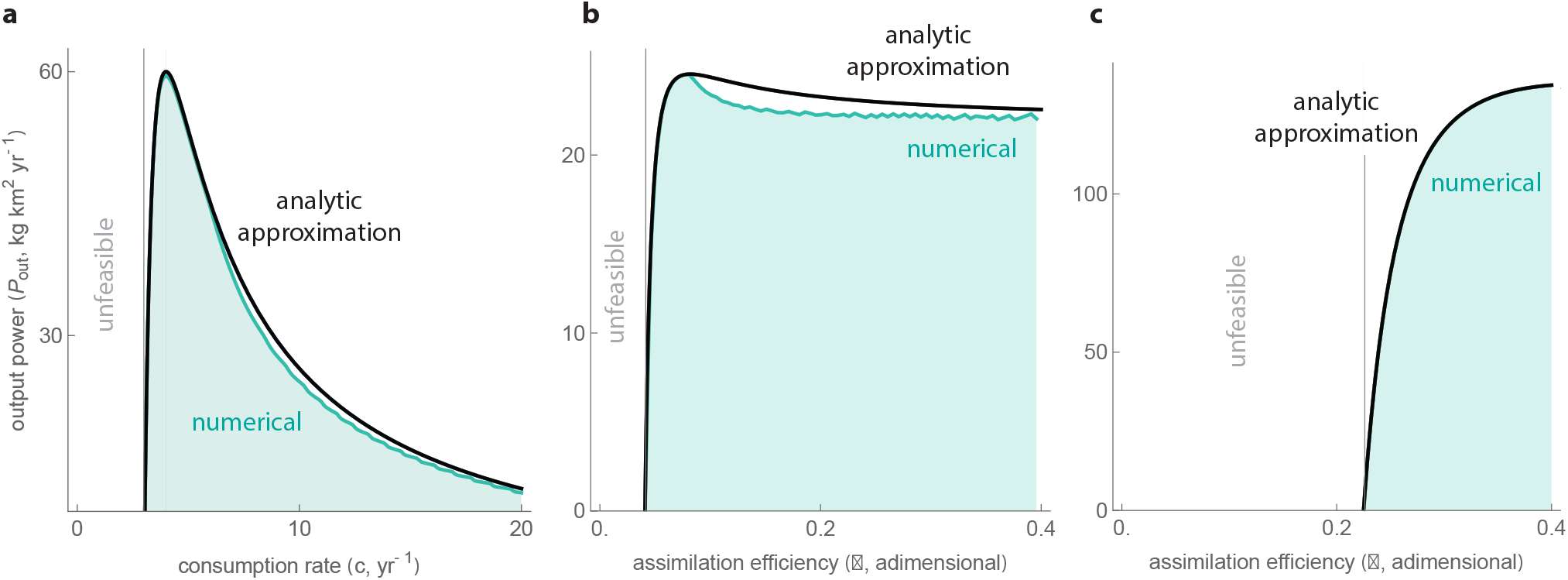
Accuracy of the approximated input and output power. Analytic approximations in (S8) (curves) versus numerical long-term averages from integrating (S1) for *T* = 1200 (points). Unless noted, parameters are as in Supplementary Table 1. (**a**) *P*_out_ versus *c* (fixed *ε*). (**b**) *P*_out_ versus *ε* (fixed *c*). (**c**) Same as (b) with a lower *c*.

### 4. Maximum output power

Consider

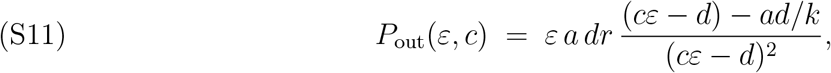

and maximize with respect to one trait subject to feasibility (S3) and *ε* ∈ [0, 1].

#### 4.1 Optimal consumption rate at fixed efficiency

Maximizing over *c* yields

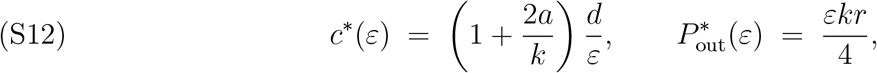

with second derivative

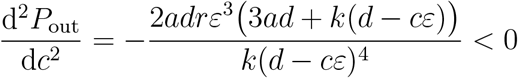

under feasibility (*cε* − *d >* 0). Note 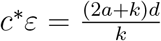, which lies strictly below the Hopf threshold (S4) for *k > a*.

#### 4.2 Optimal efficiency at fixed consumption rate

Maximizing over *ε* ∈ [0, 1] gives

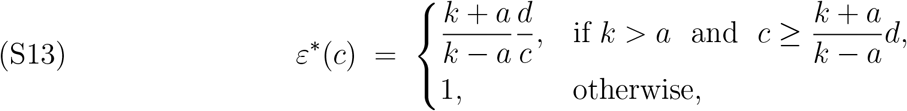

so that in the interior case

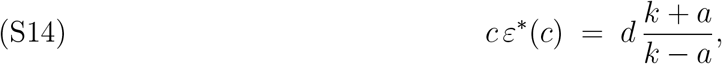

i.e., the power maximum lies exactly on the Hopf boundary (S4). When *k < a, P*_out_ increases with *ε* after feasibility and the optimum is at *ε* = 1.

**Figure S3.**
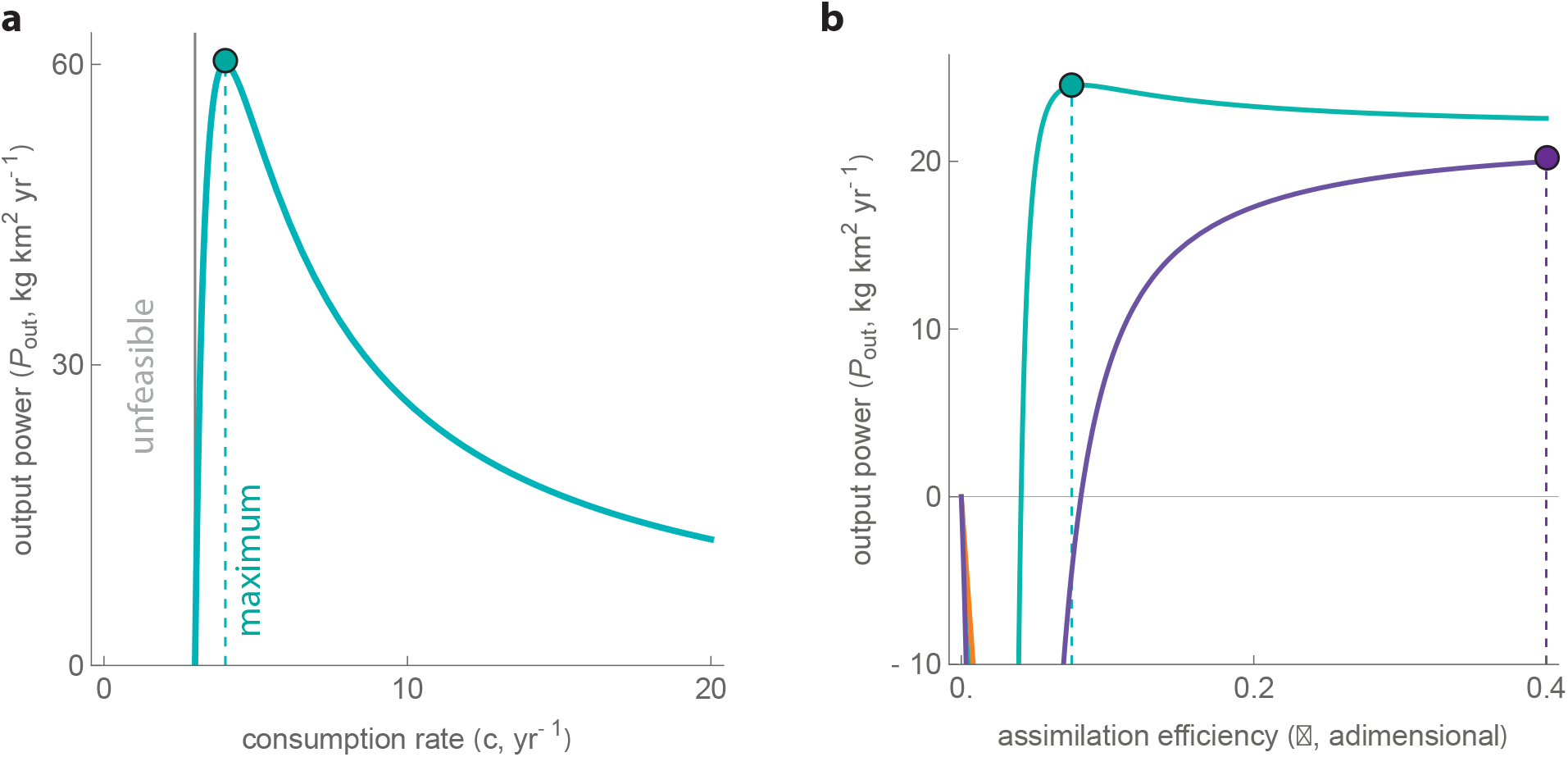
Maximum output power. Output power *P*_out_ as a function of (a) consumption rate *c* at fixed *ε* and (b) assimilation efficiency *ε* at fixed *c*. Parameters are as in Supplementary Table 1, except in panel (b) where two illustrative cases are shown: the purple curve sets *k* = *a/*2 (no interior optimum; maximum at the boundary *ε* = 1), and the orange curve sets 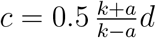 (below the interior–boundary threshold; cf. (S13)).

